# Evaluation of Various Drugs’ Influence on Alzheimer’s Disease Progress Using a New Analytical Model Based on Cellular Automata

**DOI:** 10.1101/2022.01.23.477385

**Authors:** Niloofar Jafari, Faegheh Golabi, Abbas Ebrahimi-kalan, Yashar Sarbaz

**Affiliations:** Department of Biomedical Engineering, Faculty of Advanced Medical Sciences, Tabriz University of Medical Sciences, Tabriz, Iran; Department of Neuroscience and Cognition, Faculty of Advanced Medical Sciences, Tabriz University of Medical Sciences, Tabriz, Iran; Department of Biomedical Engineering, Faculty of Electrical and Computer Engineering, University of Tabriz, Tabriz, Iran

**Keywords:** Alzheimer’s disease, Cellular Automata, Analytical Model, Drug, Amyloid-β

## Abstract

This article aims to introduce and propose a novel mathematical model for the study of Alzheimer’s disease (AD) progress. The presented model is based on Cellular Automata for better representation of AD progression. The differential equations of the Puri-Li model are utilized to calculate the number of Amyloid-β molecules. Also, a new definition for AD rate is presented in this study. Moreover, other useful factors such as Critical Rate (CR) and Warning Rate (WR) are defined to determine the status of AD progression. To get exact insight into the neuron-to-neuron communications, the model is obtained for a 3×3 neuron system to investigate the influence of drug injection on the reduction of AR, CR, and WR factors. It is shown that using drugs can decrease AR and CR factors and also enhance the WR. The presented study can be utilized for the investigation of various factors in the control and treatment of AD progression.

## 1. Introduction

Mathematical modeling is a powerful method for the forecasting, treatment, and study of various brain disorders [1, 2]. This method assists the biologists in exactly following the progress of the specific disease and also investigating the effect of various external factors such as drugs, music, exercise, and gamma-ray on the progression of brain disorders. One of the famous neurodegenerative disorders that have affected millions of people all over the world is Alzheimer’s disease (AD), which is mainly caused by the agglomeration of Amyloid-β (Aβ) molecules within the hippocampus [3].

Analytical modeling of AD has been published in many research works [4–19]. One of the famous models in this research area is the Puri-Li model which suggests seven differential rate equations to consider the cross-talk between neuron, astroglia, microglia, and Aβ. Many kinds of research are conducted based on this model, for instance, the effect of calcium Ion Hemostasis on AD progress is investigated, based on the Puri-Li model [7].

In this article, by merging the differential equation of the Puri-Li model and Cellular Automata (CA), we introduce novel analytical mathematical relations to model the neuron-to-neuron communications in a 3 × 3 neuron system. As we know, CA is one of the powerful methods which assists biologists to schematically observe and follow the progression of brain disorders. We also investigate the drugs’ effect on the reduction of AD progress, which has a large impact on AD. It is shown that the use of suitable drugs can decrease AR and CR factors and also enhance the WR.

## 2. Analytical Model

Fig.1 represents the biological structure of the neurons. As seen in Fig. 1 (a), each neuron has three main parts: cell body, axons, and dendrites. To model the neuron, as shown in Fig. 1 (b), we have ignored axon, Myelin sheath, and node of Ranvier and only kept the cell body of the neuron together with its synapses. As we know, the agglomeration of Aβ molecules occurs on synapses continuously by forming platelets. Therefore, considering the cell body of the neuron with its synapses is a realistic supposition in this study. Furthermore, the synapses have been nominated in a clockwise direction to show the formation of continuous platelets on synapses, as seen in Fig. 1 (c). We assume that each neuron has about *synapse_total_* = 1000 synapses, which means that each square synapse cell (i.e. *S*_1_,*S*_2_,*S*_3_,*S*_4_,*S*_5_,*S*_6_,*S*_7_,*S*_8_) contains *synapse_per cell_* = *synapse_total_*/8 = 125 synapses.

**Fig. 1.**
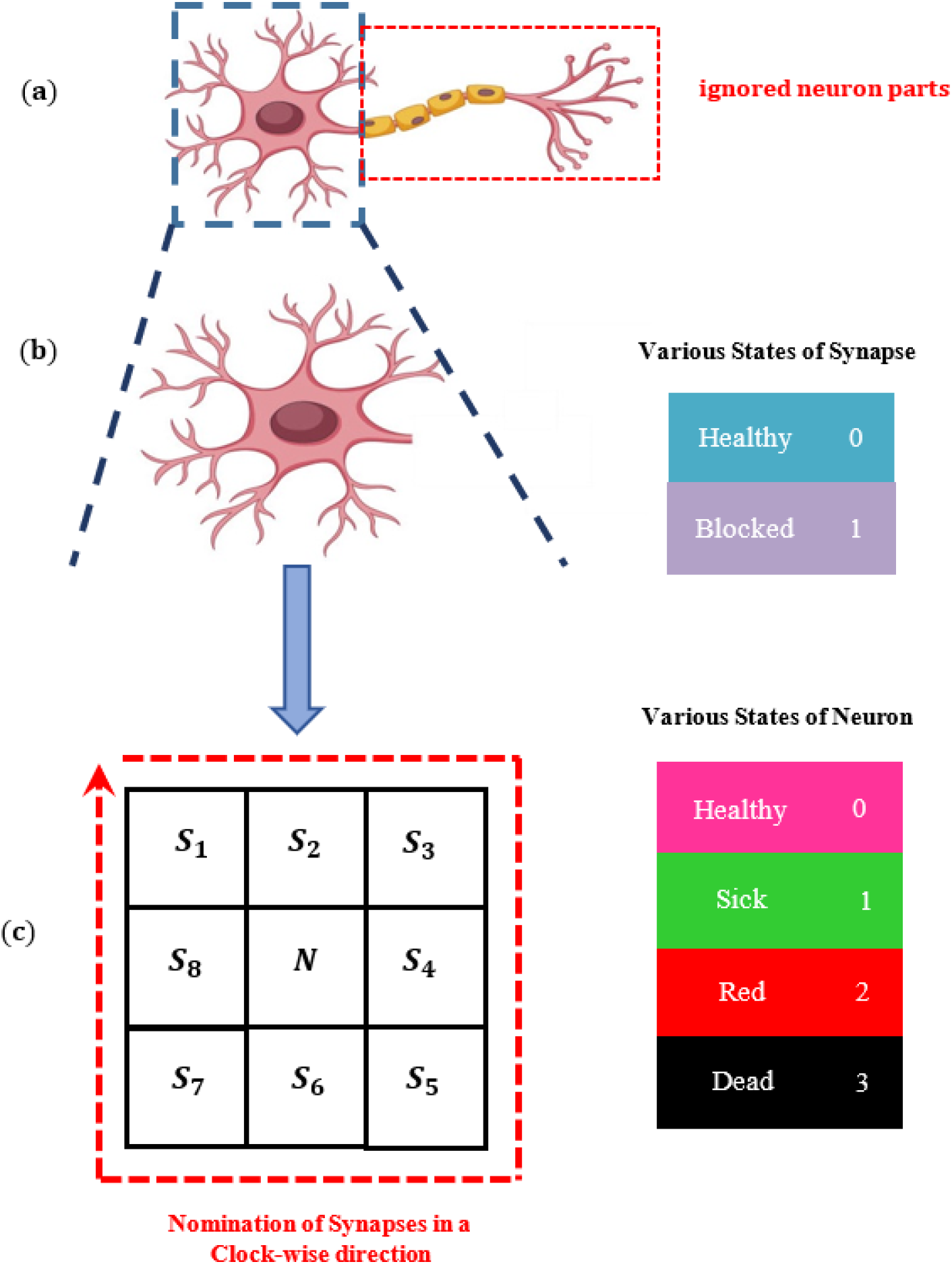
**(a)** Various parts of a neuron, including cell body, an axon, and dendrites, **(b)** The cell body (core) and its adjacent synapses, **(c)** CA Network constructed of a central cell for neuron (core) and eight cells for all its synapses. The synapse cells are named/numbered in a clockwise direction.

In our model, each synapse cell has two possible states while the central cell has 4 states. State “0” (Healthy) for a synapse cell implies a situation where the amount of Aβ molecules is not sufficient to block all synapses of the synapse cell, while state “1” (Blocked) is utilized for a synapse cell in which its whole synapses are blocked by the sufficient value of Aβ molecules. Meanwhile as illustrated in Fig. 1 (c), the neuron itself can be in one of four states including state “0” (Healthy), state “1” (sick), state “2” (red), and state “3” (dead). The healthy state implies a neuron in which its connections with other neurons are retained due to the low value of Aβ molecules and consequently low number of blocked synapse cells. For a dead neuron (state “3”), neuron-to-neuron connections are interrupted because more than 50% of the synapses have been blocked by the agglomeration of Aβ molecules. Usually, only two states (i.e., alive or dead neurons) are considered for a neuron in the literature. Since the healthy neuron gradually converts into a dead neuron, we define two new states for a neuron, i.e., sick and red states. The state “1” (sick state) is a neuron in which 25% of its synapse are blocked while the state “2” (red state) is an emergency state of a neuron in which between 25% up to 50% of its synapses are blocked. Let us suppose that the hippocampus, a small area within the brain that the AD begins, occupies about 2.5 cm^2^ in the two-dimensional surface. The number of cells is about 10^6^ by assuming that the effective area of each neuron is 250 μm^2^. This article aims to get biological insight to determine how the progression of AD occurs and the simulation of these huge numbers is outside the scope of this article. Therefore, we consider a 3 × 3 neuron network here, as depicted in Fig. 2.

**Fig. 2.**
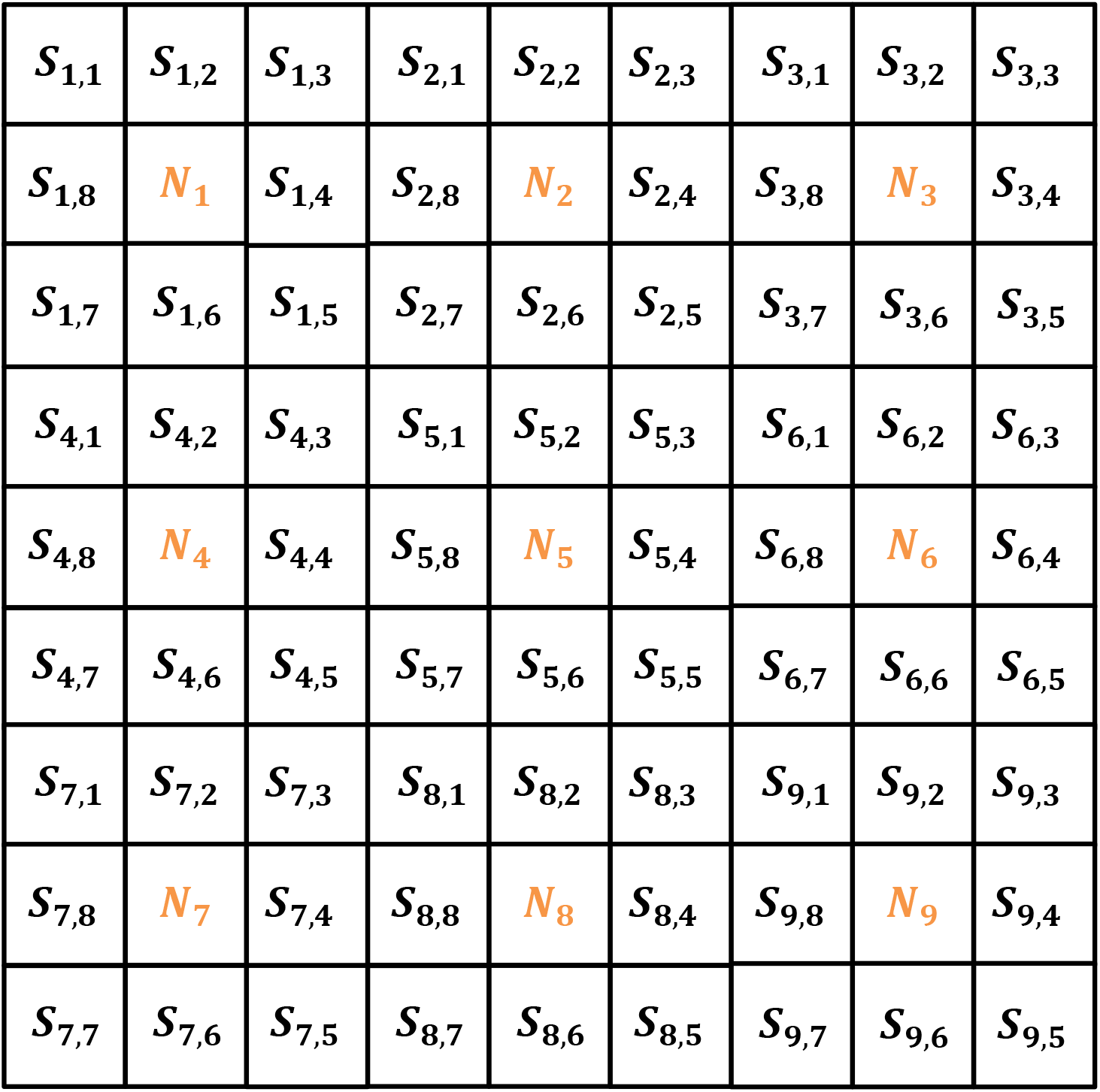
Cellular automata for 3 × 3– neuron network.

As explained before, the main parameter in AD is the amount of Aβ molecules, which can be calculated by the Puri-Li dynamic model for each neuron in a 3 × 3 network [4]:

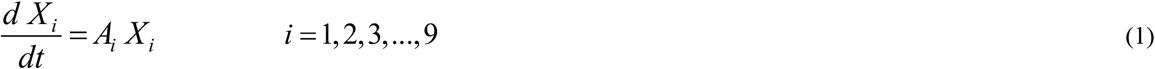

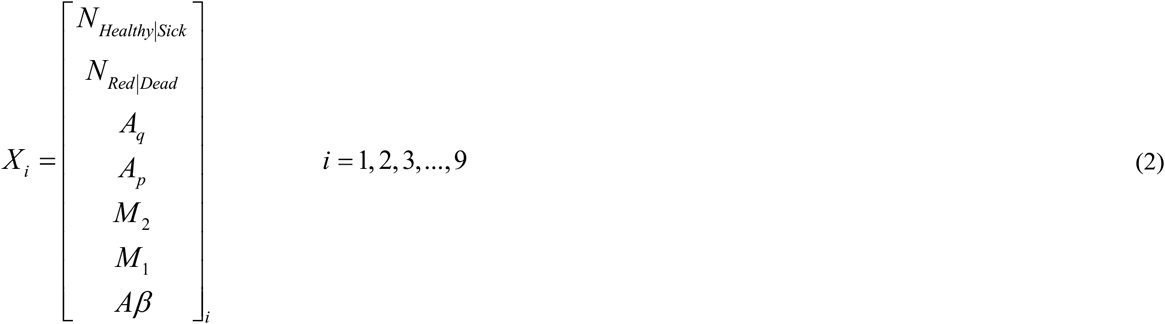

where

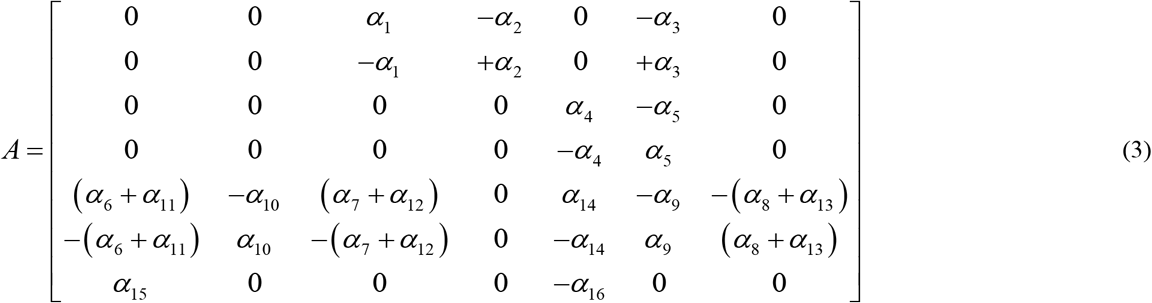

is a matrix, as defined by Puri and Li [4]. In (2), *A_q_,A_p_, M_2_, M_1_, Aβ* are the number of quiescent astroglia, proliferating astroglia, anti-inflammatory state, pro-inflammatory state, and Aβ molecules, respectively [4]. In the general form, one can express the response of (1-3) for *i*-th neuron (the primary values of *x*_0,*i*_) as follows:

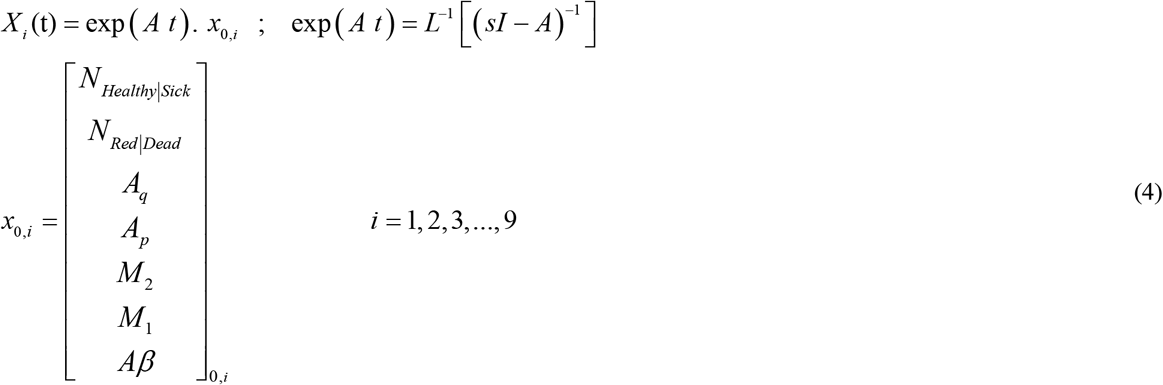

In equation (4), *Aβ* determines the whole value of Aβ molecules for all synapses of the *i*-th neuron. Now, the number of blocked synapse cells at *t* = *t*_0_ is obtained:

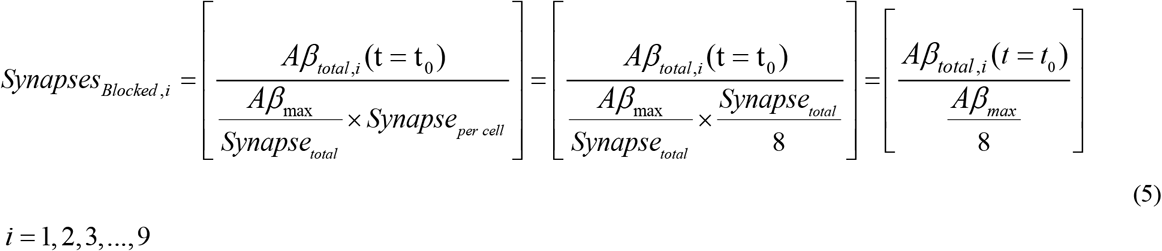

Based on equation (5), *Aβ_total,N_*(*t* = *t*_0_) and *Aβ_max_* are the value of Aβ molecules (calculated by the Puri-Li model) and the maximum value of Aβ molecules to be blocked completely, respectively. Moreover, the “[]” is used for the *floor* function. Now, the synapse states for the *i*-th neuron can be derived:

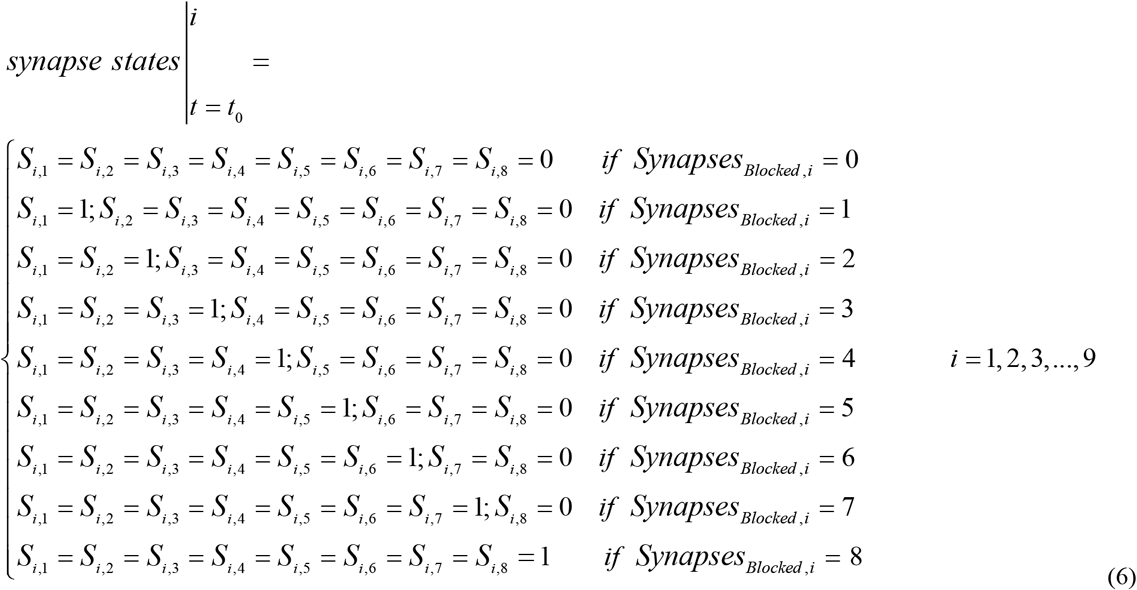

It should be explained that the presented formula in (5) cannot model the neuron-to-neuron communications effectively since it only calculates the number of self-blocked synapse cells. Within the brain, the synapses of each neuron send/receive signals to/from adjacent and non-adjacent neurons. Here, neuron-to-neuron communication of adjacent neurons is modeled for a 3 × 3-neuron network:

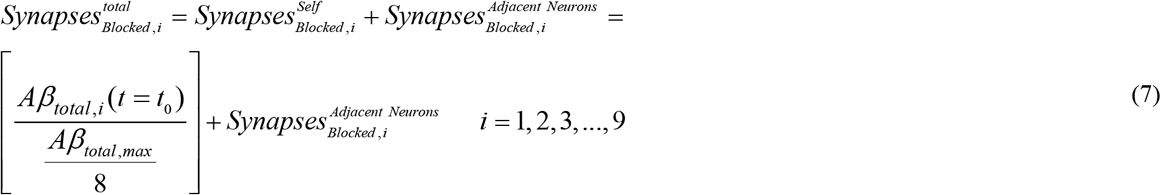

As shown in Fig. 3, in our model of 3 × 3– Neuron Network, three kinds of neurons exist: *1-Corner Neurons* (i.e. *N*_1_, *N*_3_, *N*_7_, *N*_9_, their communications are illustrated by green color), *2-Row and Column Neurons* (i.e. *N*_2_, *N*_4_, *N*_6_, *N*_8_, their communications are shown by blue color), and *3-Central Neuron* (i.e. *N*_5_, it has been highlighted by brown color). The neuron state for each kind of these categories can be obtained as follows:

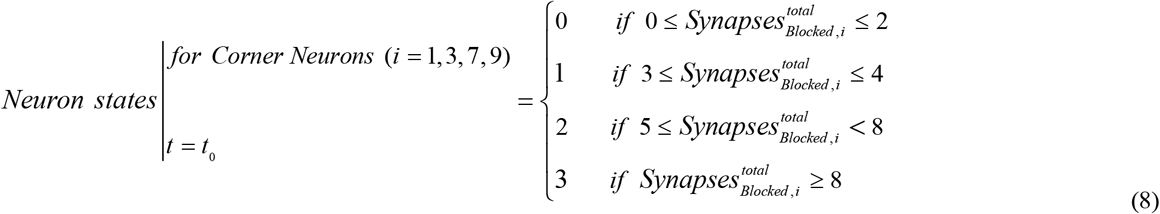

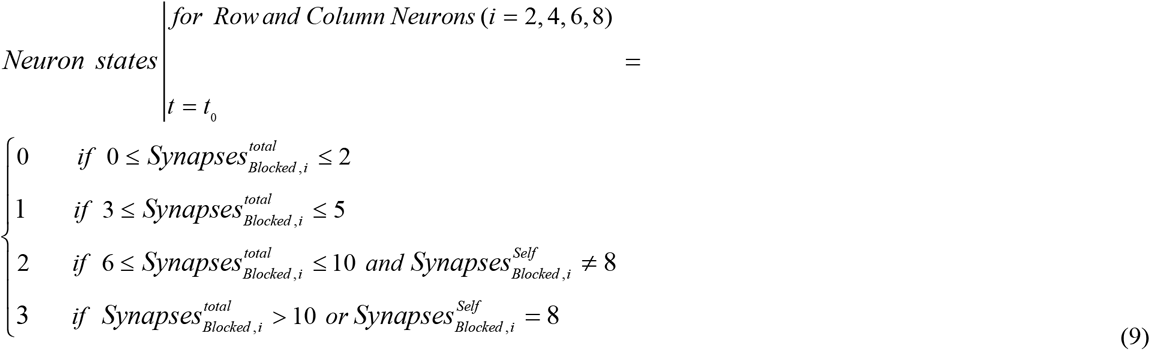

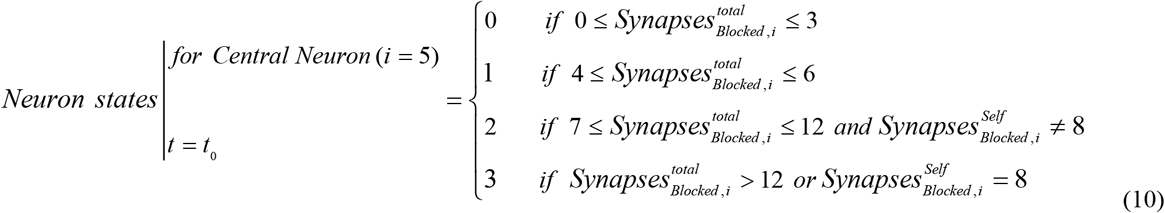

**Fig. 3.**
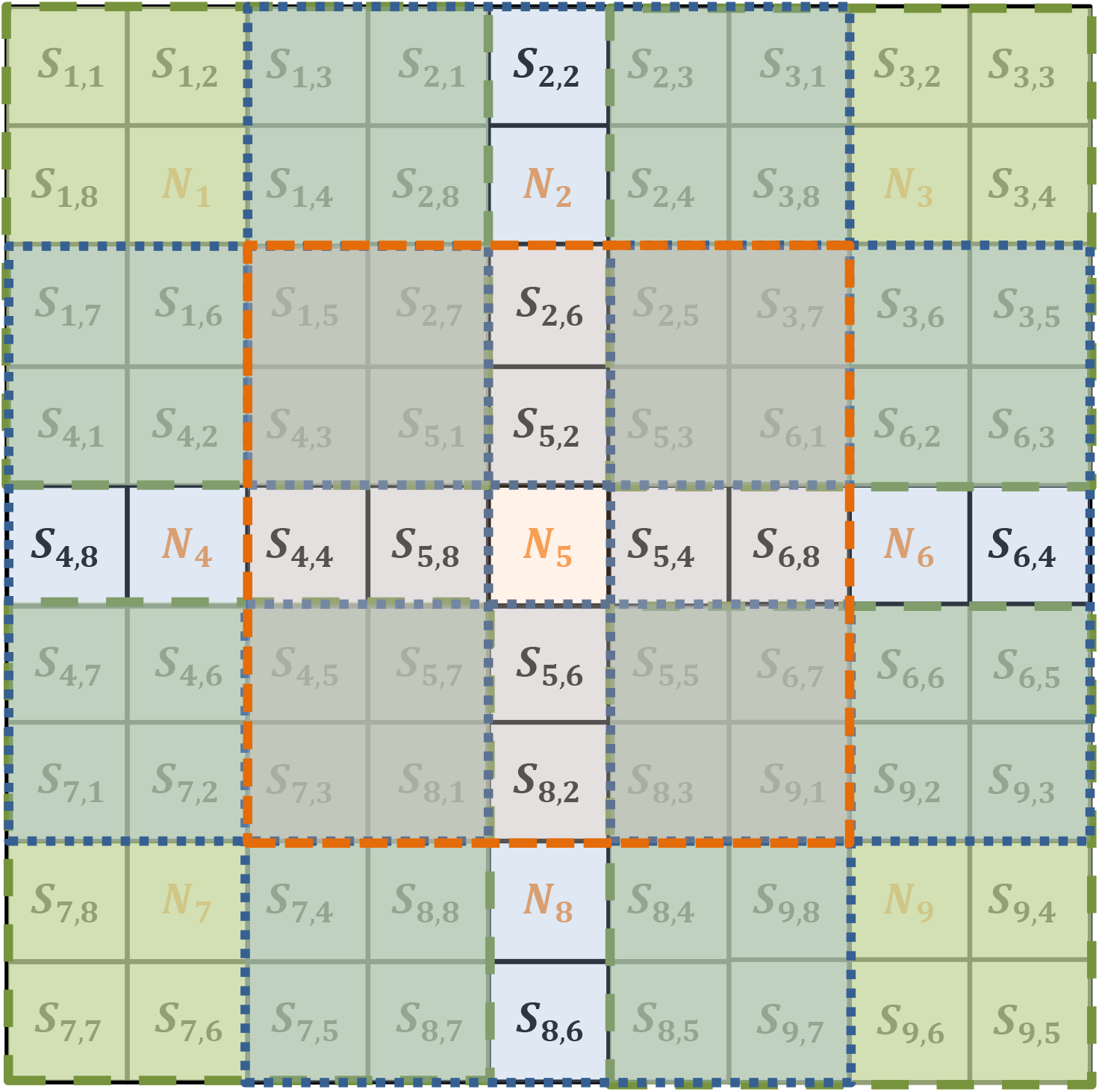
Neuron-to-neuron communications in 3 × 3– neuron network. Corner neurons, row, and column neurons, and central neurons are illustrated by green, blue, and brown colors, respectively.

To effectively control the individual status, the following parameters are defined:

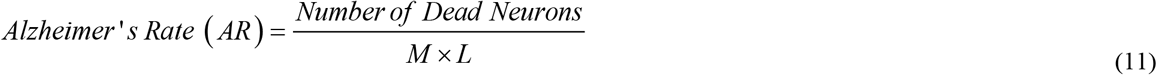

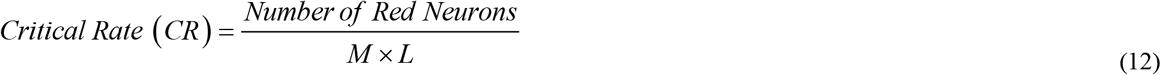

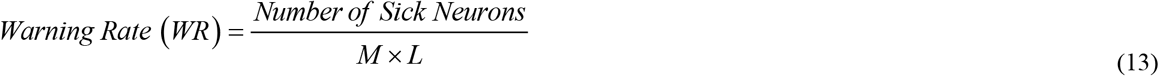

By supposing the time step of *t* = *t*_0_, the synapse state for the next time step, i.e., *t* = 2*t*_0_, can be obtained by using Puri-Li equations:

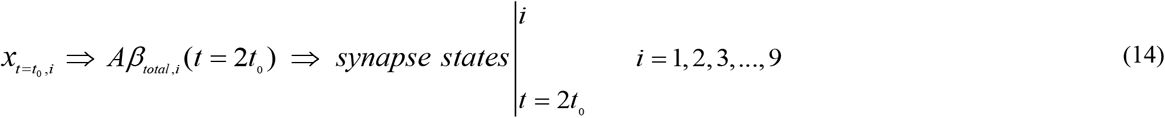

However, some internal mechanisms within the brain and also some external factors such as drugs, can modify the amount of Aβ molecules. Therefore, the primary values for *i*-th neuron at *t* = *t*_0_ can be updated as:

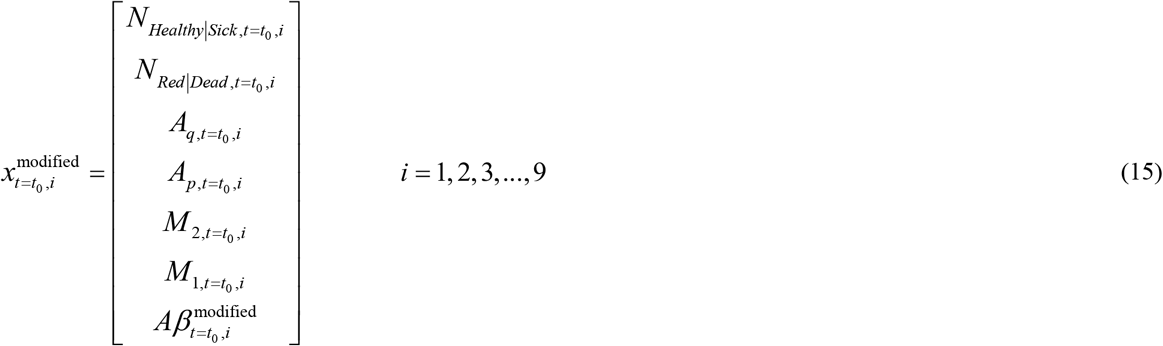

where 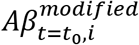 in (15) is defined as:

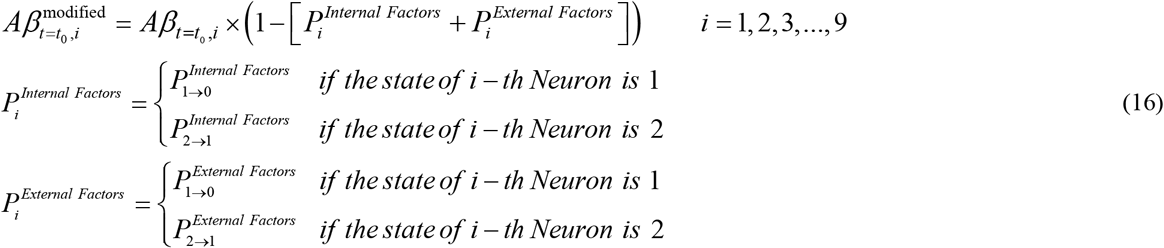

For instance, 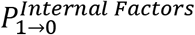 is the probability of the state of the *i*-th neuron changing from the sick state (state “1”) to the healthy state (state “0”) as result of internal factors.

## 3. Results and Discussions

This section studies the numerical results of the proposed model for a 3 ×3 neuron network coded by software. The simulation parameters of the CA model and the mathematical parameters of the Puri-Li model are given in Table. 1 and Table. 2, respectively. In Table. 3, the probabilistic functions for three AD drugs, i.e., Rivastigmine, Donepezil, and Galantamine, are given. Moreover, the initial values of the neuron parameters have been given in Table. 4.

**Table 1.**
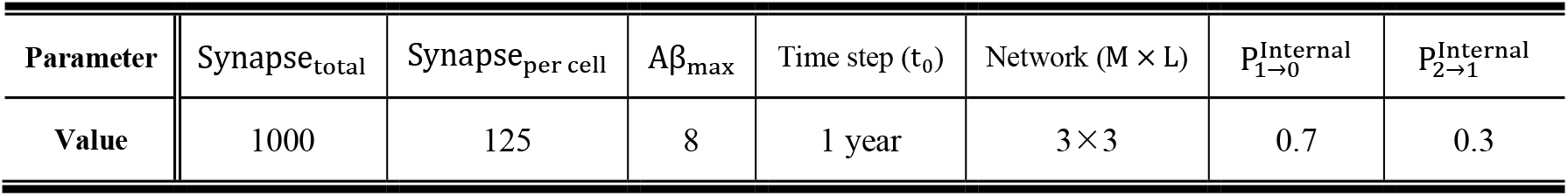
Simulation Parameters of CA Network.

**Table 2.** Mathematical Parameters of Puri-Li model (1/*year*).

| Parameter | $\alpha_1$ | $\alpha_2$ | $\alpha_3$ | $\alpha_4$ | $\alpha_5$ | $\alpha_6$ | $\alpha_7$ | $\alpha_8$ | $\alpha_9$ | $\alpha_{10}$ | $\alpha_{11}$ | $\alpha_{12}$ | $\alpha_{13}$ | $\alpha_{14}$ | $\alpha_{15}$ | $\alpha_{16}$ |
| --- | --- | --- | --- | --- | --- | --- | --- | --- | --- | --- | --- | --- | --- | --- | --- | --- |
| Value | $10^{-5}$ | $10^{-3}$ | $10^{-2}$ | $10^{-4}$ | $10^{-2}$ | $10^{-2}$ | $10^{-4}$ | $10^{-2}$ | $10^{-2}$ | $10^{-2}$ | $10^{-2}$ | $10^{-4}$ | $10^{-2}$ | $10^{-4}$ | 1 | $10^{-2}$ |

**Table 3.** Probabilistic functions of various drugs.

| Rivastigmine |  | Donepezil |  | Galantamine |  |
| --- | --- | --- | --- | --- | --- |
| $p_{1 \rightarrow 0}^{\text{External}}$ | $p_{2 \rightarrow 1}^{\text{External}}$ | $p_{1 \rightarrow 0}^{\text{External}}$ | $p_{2 \rightarrow 1}^{\text{External}}$ | $p_{1 \rightarrow 0}^{\text{External}}$ | $p_{2 \rightarrow 1}^{\text{External}}$ |
| 0.08 | 0.07 | 0.19 | 0.17 | 0.16 | 0.15 |

**Table 4.**
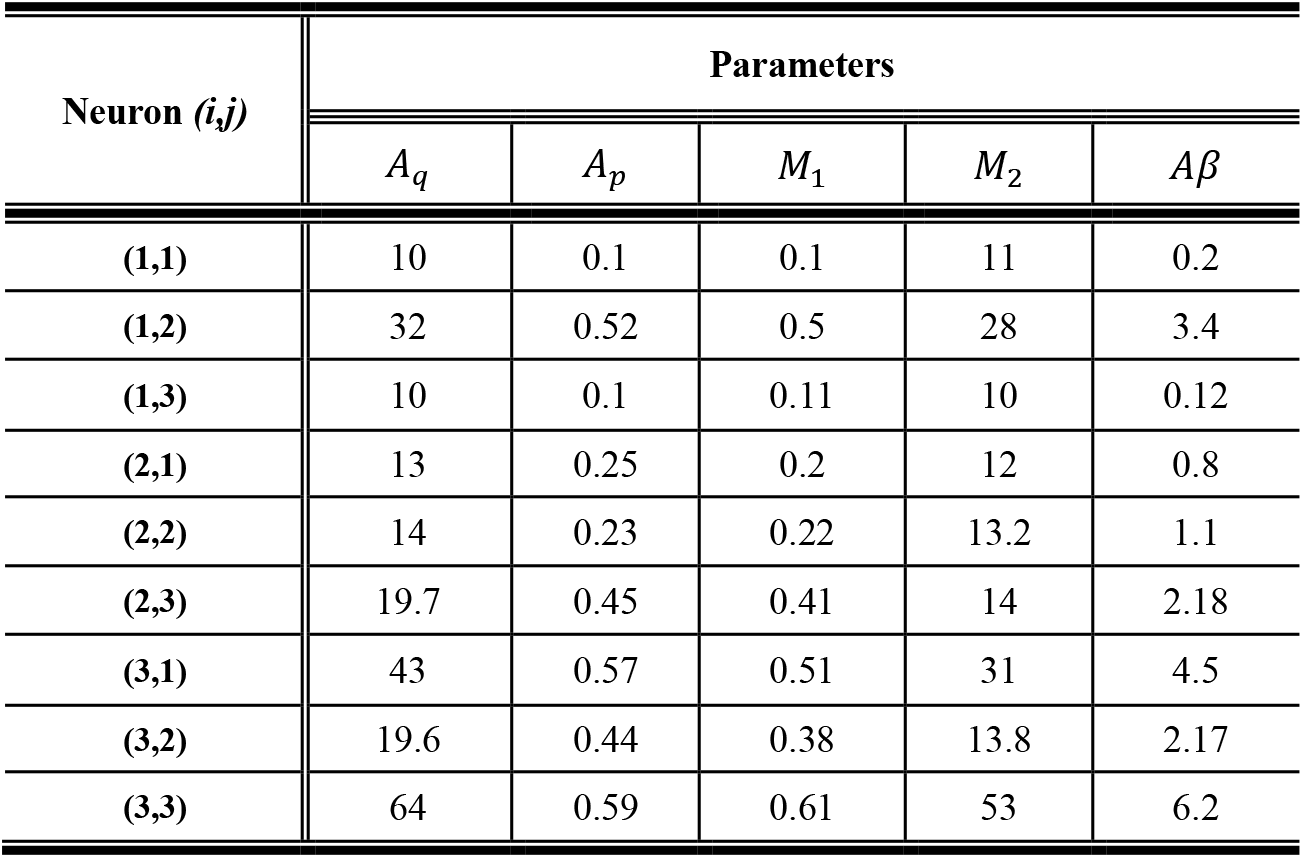
Initial values of various parameters of neurons in a 3 × 3 network.

| Neuron ($i,j$) | Parameters | | | | |
| --- | --- | --- | --- | --- | --- |
| | $A_q$ | $A_p$ | $M_1$ | $M_2$ | $A\beta$ |
| (1,1) | 10 | 0.1 | 0.1 | 11 | 0.2 |
| (1,2) | 32 | 0.52 | 0.5 | 28 | 3.4 |
| (1,3) | 10 | 0.1 | 0.11 | 10 | 0.12 |
| (2,1) | 13 | 0.25 | 0.2 | 12 | 0.8 |
| (2,2) | 14 | 0.23 | 0.22 | 13.2 | 1.1 |
| (2,3) | 19.7 | 0.45 | 0.41 | 14 | 2.18 |
| (3,1) | 43 | 0.57 | 0.51 | 31 | 4.5 |
| (3,2) | 19.6 | 0.44 | 0.38 | 13.8 | 2.17 |
| (3,3) | 64 | 0.59 | 0.61 | 53 | 6.2 |

In Fig. 4, the CA representation of neurons and their synapses for a sample 3× 3 neuron network has been depicted at *t* = 0 and *t* = 1 *year*. As mentioned previously, the status of each neuron depends on the states of its synapses and the synapses of adjacent neurons. For instance, all synapses of neuron (1,3) are healthy in Fig. 4 (a) while its status is sick, due to the blocked synapse cells of its adjacent neurons (i.e., neuron (1,2) and neuron (2,3)). After one year, although AR is still zero, CR reaches 88.8%, which is a critical situation. To control this critical situation, by reducing the CR factor, some internal mechanisms within the hippocampus can help to decrease the amount of Aβ. These mechanisms are essential but not adequate. Therefore, an effective external factor should be added to this system to reduce CR. One practical way is the usage of drugs such as Rivastigmine, Donepezil, and Galantamine. Here, we aim to investigate the influence of these drugs on the enhancement of critical situations in a 3 × 3 neuron network.

**Fig. 4.**
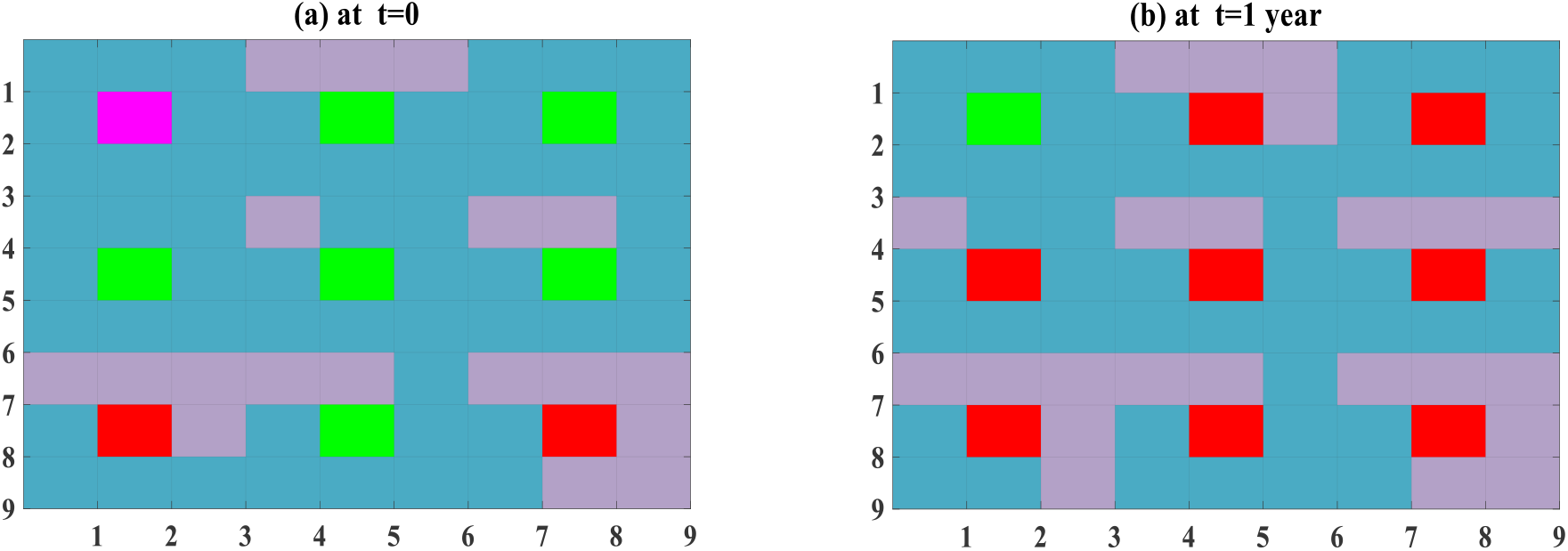
Initial states of neurons and their synapses in CA model of sample 3 × 3 network at: **(a)** *t* = 0, **(b)** *t* = 1 *year*.

Fig. 5 shows the analytical results of CA in sample 3×3 neuron network after two years, without any drugs and with the Rivastigmine injection. One can observe from Fig. 5 (a) that internal mechanisms decrease CR (*CR* ≈ 11.1 %) in the second year but the WR has reached *WR ≈* 88.8 % meaning that the system is already is in an emergency state. As shown in Fig. 5 (b), by injecting Rivastigmine during the second year, CR becomes zero and also the numbers of blocked synapses reduce. As a result, the Rivastigmine injection converts the critical state into a warning state, which is desirable.

**Fig. 5.**
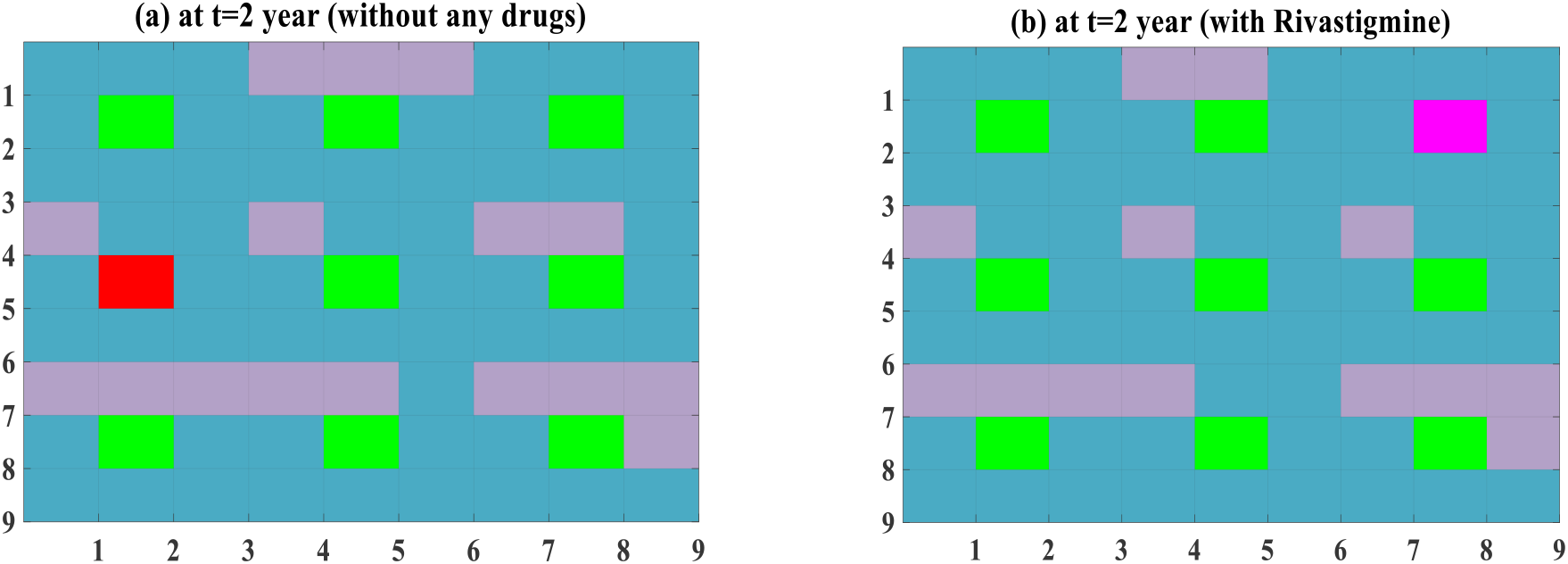
CA model of neurons and their synapses in sample 3 × 3 network at *t* = 2 *years*: **(a)** without any drugs, **(b)** with Rivastigmine.

To decrease the WR factor, a strong drug such as Galantamine can also be utilized. In Fig. 6, the analytical results of CA in a 3×3 neuron network after two years, with and without the Galantamine injection, are depicted. One can see from Fig. 6 (b) that *CR = 0* is similar to Rivastigmine injection. However, Galantamine has reduced the WR factor (*WR* ≈ 66.6 %), which is better than Rivastigmine injection.

**Fig. 6.**
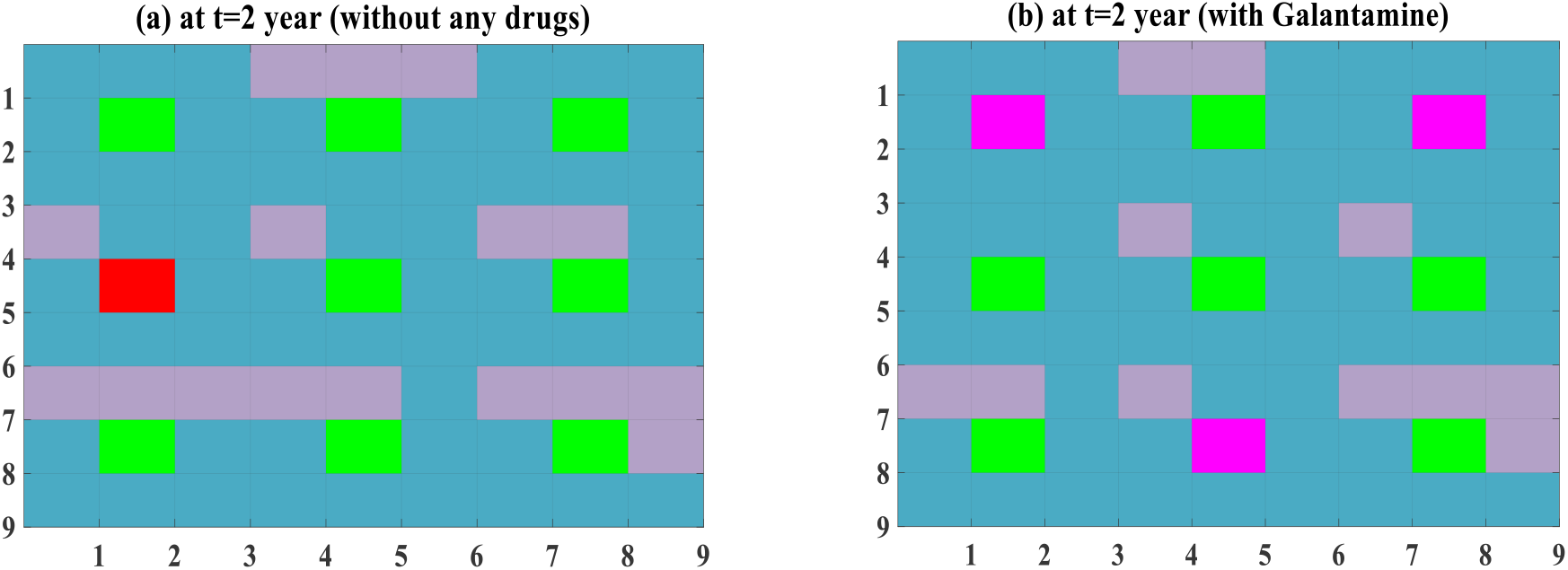
CA model of neurons and their synapses in sample 3 × 3 network at *t* = 2 *years*: **(a)** without any drugs, **(b)** with Galantamine.

As a final point, we studied the effect of Donepezil on the analytical results of CA in a 3 × 3 neuron system after two years, as shown in Fig. 7. It is clear from this figure that the CR factor reduces to zero by Donepezil Injection. Compared to other drugs i.e., Rivastigmine and Galantamine, the WR factor reaches *WR* ≈ 55.5 %, which is a great enhancement. Hence, utilizing a drug such as Donepezil can convert the emergency state into a warning state, and also give a lower WR factor in which the neurons’ states are very stable in this situation.

**Fig. 7.**
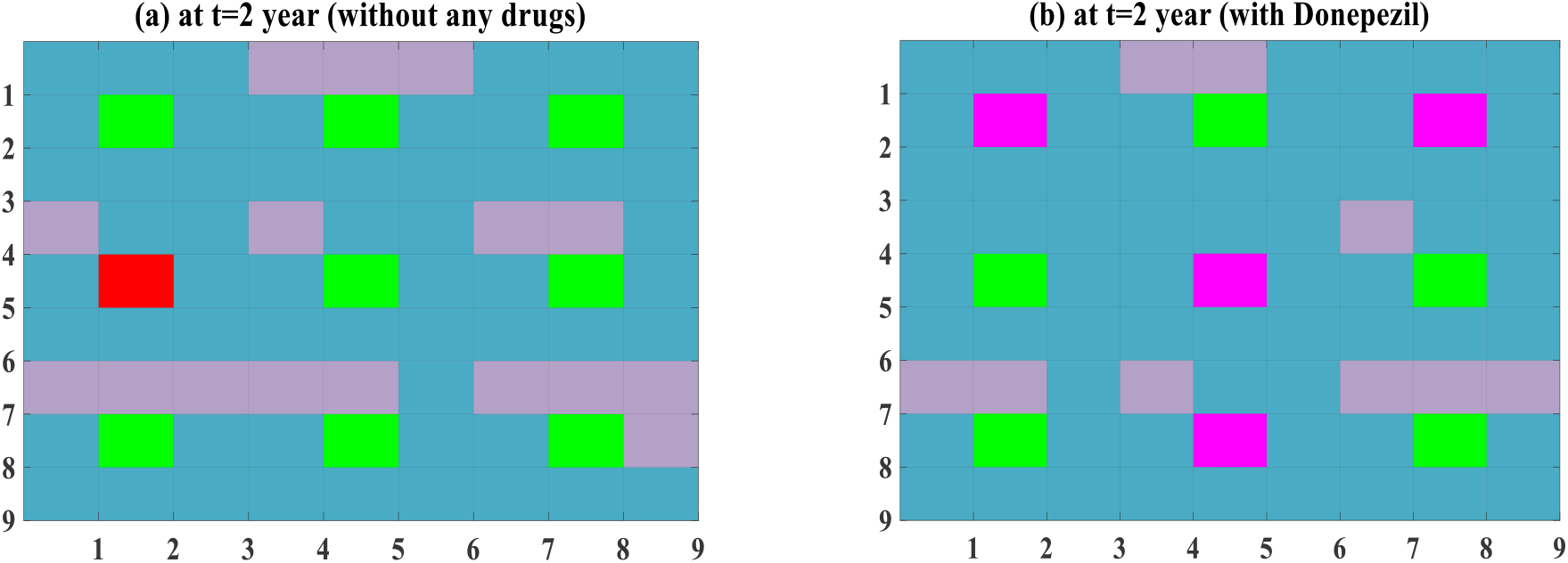
CA model of neurons and their synapses in sample 3 × 3 network at *t =* 2 *years*: **(a)** without any drugs, **(b)** with Donepezil.

The diagrams in Figures (8) to (10) show the effect of drugs on the critical and warning rate. The use of drugs improves the critical rate and warning rate in Alzheimer’s disease.

**Fig. 8.**
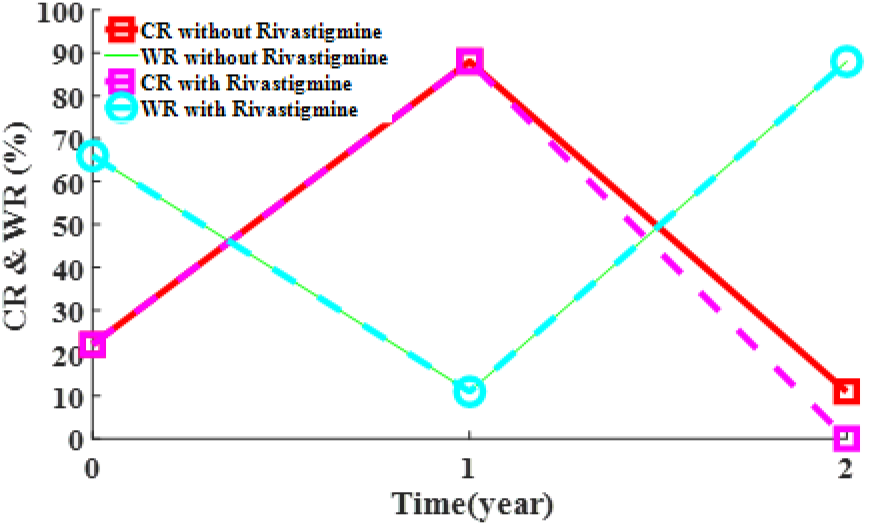
The effect of Rivastigmine on critical and warning rate.

**Fig. 9.**
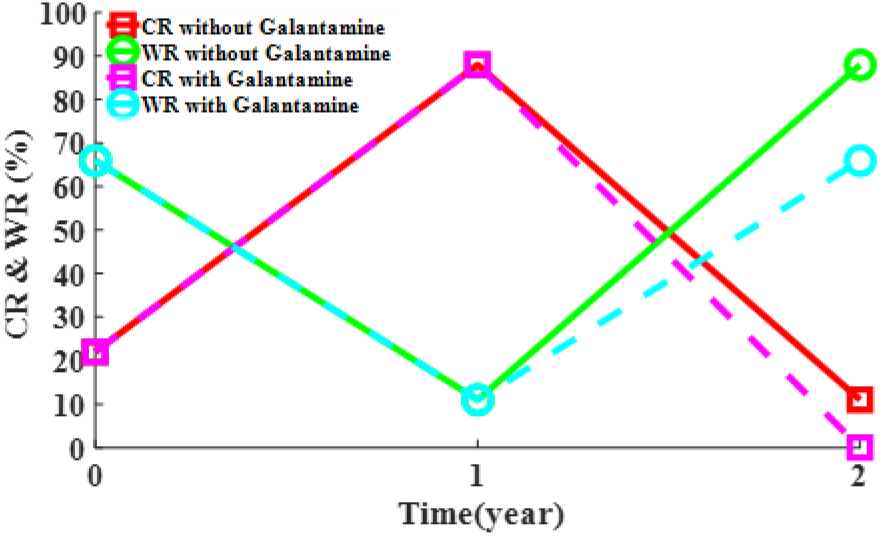
The effect of Galantamine on critical and warning rate.

**Fig. 10.**
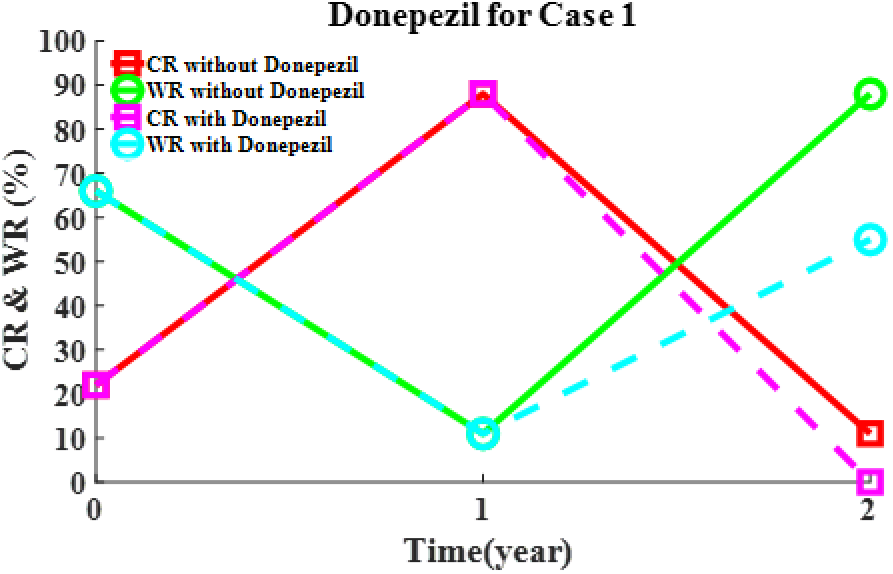
The effect of Donepezil on the critical and warning rate.

## 4. Conclusion

In this paper, we proposed a new model for the study of drug injection on AD progress. This model was presented by the compounding of Puri-Li equations in the Cellular Automata. As a special case of general neuron network, a 3 × 3 neuron network was considered and the states of neurons and synapse cells were schematically shown. New definitions for AR, CR, and WR factors were presented to determine the stage of AD. It was shown that the drug injection can decrease the AR and CR factors to zero. Besides, choosing a strong drug such as Donepezil could reduce the WR factor, which is desirable. The proposed model can also be utilized for the study and forecasting the effect of various factors such as exercise, music, drugs, Gamma-ray, etc. on the AD progress.

## Declarations

### Ethical statement

This work is partially supported by Vice-Chancellor in Research Affair-Tabriz University of Medical Sciences under Ethical Code No. IR.TBZMED.VCR. REC.1399.377. The funders had no role in study design, analysis, numerical simulations, the decision to publish, or preparation of the manuscript.

